# Computational modeling of methylation impact of AML drivers reveals new pathways and refines AML risk-stratification

**DOI:** 10.1101/2023.05.17.541249

**Authors:** Burcu Gurun, Jeffrey W. Tyner, Emek Demir, Brian J. Druker, Paul T. Spellman

## Abstract

Decades before its clinical onset, epigenetic changes start to accumulate in the progenitor cells of Acute Myelogenous Leukemia (AML). Delineating these changes can improve risk-stratification for patients and shed insights into AML etiology, dynamics and mechanisms. Towards this goal, we extracted “epigenetic signatures” through two parallel machine learning approaches: a supervised regression model using frequently mutated genes as labels and an unsupervised topic modeling approach to factorize covarying epigenetic changes into a small number of “topics”. First, we created regression models for *DNMT3A* and *TET2*, the two most frequently mutated epigenetic drivers in AML. Our model differentiated wild-type vs. mutant genotypes based on their downstream epigenetic impacts with very high accuracy: AUROC 0.9 and 0.8, respectively. Methylation loci frequently selected by the models recapitulated known downstream pathways and identified several novel recurrent targets. Second, we used topic modeling to systematically factorize the high dimensional methylation profiles to a latent space of 15 topics. We annotated identified topics with biological and clinical features such as mutation status, prior malignancy and ELN criteria. Topic modeling successfully deconvoluted the combined effects of multiple upstream epigenetic drivers into individual topics including relatively infrequent cytogenetic events, improving the methylation-based subtyping of AML. Furthermore, they revealed complimentary and synergistic interactions between drivers, grouped them based on the similarity of their downstream methylation impact and linked them to prognostic criteria. Our models identify new signatures and methylation pathways, refine risk-stratification and inform detection and drug response studies for AML patients.

**KEY POINTS:**

- Supervised and unsupervised models reveal new methylation pathways of AML driver events and validate previously known associations.
- Individual *DNMT3A* and *TET2* signatures are accurate and robust, despite the complex genetic and epigenetic make-up of samples at diagnosis.
- Unsupervised topic modeling factorizes covarying methylation changes and isolates methylation signatures caused by rare mutations.
- Topic modeling reveals a group of mutations with similar downstream methylation impacts and mapped to adverse-risk class by ELN.
- Topic modeling uncovers methylation signatures of infrequent cytogenetic events, significantly improving methylation-based subtyping.
- Our models can be leveraged to build predictive models for AML-risk.
- Our models show that cytogenetic events, such as t(15;17) have widespread *trans* downstream methylation impacts.

## INTRODUCTION

In many Acute Myeloid Leukemia (AML) cases, precancerous clonal expansions start in healthy tissue decades before the cancer onset^1–3^. A hematopoietic stem cell may follow different paths to AML gaining the necessary pathological features (e.g., increased proliferation, abnormal morphology, and impaired differentiation of myeloid lineage cells, or combinations thereof) at different rates,^4^ dictated by the interplay between the type and order of mutations, epigenetic events and tumor micro-environment^5^.

Before the acute presentation of a myeloid malignancy with anemia, neutropenia and thrombocytopenia, the disease is clinically silent and hard to detect. Recent single-cell sequencing studies^6,7^ showed that this early phase is primarily driven by a small set of mutations^8–10^. Clones frequently harbor multiple co-occurring mutations including *DNMT3A, TET2* and *ASXL1*, which are epigenetic regulators that change the normal physiological methylation landscape when mutated ^6,7,11^, thus conferring a broad risk of progression to a myeloid malignancy with the acquisition of a cooperating mutation^12^. Once the disease progresses to an advanced stage it develops substantial clonal heterogeneity and becomes difficult to treat; only 20% of the patients survive for more than 5 years after diagnosis and recurrence is common even after complete remission^13^.

These early drivers lead to a large number of Differentially Methylated Regions (DMRs) in the genome^14^. Their effects can be captured as distinct “signatures” to gain a better understanding of the mechanisms and dynamics that drive early disease. DMR signatures could be used for characterizing aberrant methylation in cancers^15^, tumor sub-classification and distinguishing the tissue of tumor origin^16–18^. For example, Vosberg *et al*. ^16^ showed that the AML clinical risk stratification based on genetic mutations European LeukemiaNet (ELN-2017) has good concordance with the DNA methylation based clustering. The authors suggested that the DNA methylation profiling could be used for AML risk stratification as subgroups of epigenetically homogeneous AML patients differ significantly in clinical outcomes. Similarly, Cabezon *et al*.^19^ demonstrated that different methylation signatures at the time of diagnosis could predict response to azacytidine, a hypomethylating agent.

Compared to genomic profiles, methylome profiles offer two key benefits for liquid biopsy applications. First, a single genomic alteration can be associated with thousands of methylation changes, leading to significant signal diversity if the methylation changes have selective advantage. Secondly, various low frequency genomic alterations that have similar downstream effects can be grouped into methylation factors. These advantages are critical in both cancer early detection and disease monitoring applications, as they increase the power to detect rare subclonal expansions of pre-malignant diseases.

In this study we investigated how well the upstream genomic events are reflected in the methylome, and whether common downstream effects of groups of genomic events can be detected. To achieve this objective, we systematically modeled methylation signatures in AML using methylation, genetic, and clinical profiling data from 220 AML patients in the BeatAML cohort^20^. We used two statistical approaches to capture these signatures. First, we identified frequent epigenetic signatures linked to mutations in *DNMT3A* and *TET2* genes and used supervised regression models to deduce methylation activities. We demonstrated that epigenetic signatures in methylation pathways were associated with *DNMT3A* and *TET2* mutations (**Figure 1**). Next, we utilized unsupervised topic modeling^21,22^ to develop a latent representation of the methylation landscape and labeled methylation signatures using biological and clinical factors. We showed that topic modeling can accurately identify the impacts of all major drivers, and these signatures are highly correlated with ELN-2017^23^. Our findings suggest that our approach can significantly improve risk-stratification for AML patients by revealing the previously unknown methylation signatures of relatively rare cytogenetic events, which will also greatly improve the subsequent methylation-based subtyping of patients for drug response studies and provide a framework for the usage of methylation as a biomarker.

**Figure 1:**
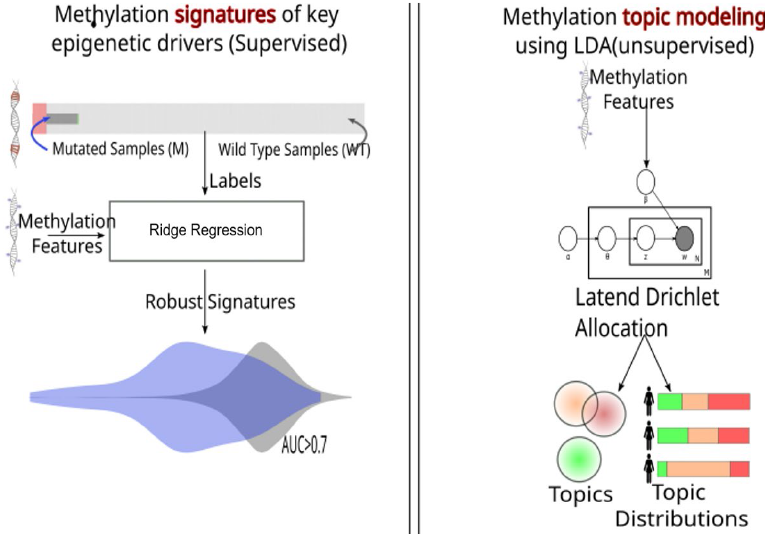
Overall Approach: Inferring methylation signatures that drives AML from methylation profiles: We use two parallel approaches: (a) regression models of epigenetic impact of epigenetic regulator mutations. (b) a latent representation of the epigenetic landscape using topic modeling.

## METHODS

### Dataset

We used a subset of the BeatAML cohort that consists of 220 post-diagnosis AML samples profiled with three modalities: 1) Illumina Infinium MethylationEPIC assay^24^ 2) matching exome sequencing and cytogenetic information and 3) matching clinical profiling^20^.

Methylation values of each probe was normalized to follow an approximate Beta – valued distribution, with constrained to lie between 0 (unmethylated locus) and 1 (methylated locus), normalized based on the BMIQ method^25^. Beta values indicate the probability that the corresponding locus is methylated.

### Approach for supervised models

We used an Elastic Net based approach^26^ to infer wild type vs mutant genotypes for *DNMT3A* and *TET2* based on their downstream epigenetic impacts. We trained regression models with different regularization parameters and evaluated statistical power and sample bias with bootstrapping. Samples were divided into two sets: training and validation (80%), and test (20%). Training and validation set were used to optimize hyperparameter(s) with 10-fold cross validation. The test set was only used for obtaining performance measures of the model on new data. We used a variance filter of top 1% most variable methylation sites. We obtained the performance measures using receiver operator characteristic (ROC) curve analysis from the ROCR package^27^. We evaluated 25 different 80% training and 20% test sets splits and observed the standard deviation of resulting models as well as hyperparameters to ensure that the models are highly robust to sampling bias. For further validation, we evaluated the biological relevance of the results by extracting the loci with the highest feature (CpG sites) coefficients and annotated them. For annotation purposes, we used ridge regularized models to have more stable feature weights. In ridge regression, all coefficients are assigned with a nonzero value during regularization and coefficients with high absolute values are important in terms of predicting the mutation status, positive coefficients specifying hypermethylation and negative values specifying hypomethylation. We extracted a list of loci and annotated the genome based on their closest gene on the genome to reveal the relationship between the selected most important chromosome sites and the epigenetic regulator under investigation.

### Approach for unsupervised models

We use topic modeling to reduce the feature space of 866,800 methylation loci to a latent space of small number of topics (factors) and represent each sample as a combination of a few topics. We used a Latent Dirichlet Allocation (LDA)^28^ based approach implemented in the topicmodels package^29^. This implementation provides explicit quantification of uncertainty, important for evaluating and comparing topics. While implementing the topic modeling, we optimized the model parameters including topic size as well as the size of the chromosome sites/probes provided as an input to the model. For validation with topic perplexity, the model evaluates how well the input matrix (methylation values for all chromosome sites of a new patient) is reconstructed from the output matrices, finding the hyperparameters minimizing the reconstruction error iteratively. We further evaluated model performance using topic coherence with biological and clinical information mapping. After inferring topics and assigning topic values to each patient with the specified optimal topic size of 15 and by taking the top 1% of the most variable genomic sites across patients, we annotated the topics with biological and clinical information. Then, we systematically tabulated the statistical enrichment of known factors for each topic using –log p-values based on the student t-test.

### Data sharing statement

All data used were from previously published studies^20,24^.

The source code for the analysis is available at https://github.com/gurundem/AML_methylation_signatures

## RESULTS

### *DNMT3A* and *TET2* have robust downstream methylation signatures

We computationally constructed signatures for the downstream impact of *DNMT3A* and *TET2*, and then, we used these signatures to shed light on their methylation pathways in AML.

Our rationale for modeling *DNMT3A* and *TET2* mutations has been four-fold: 1) For a given gene, our regression models require sufficient number of mutant cases in a given cohort to produce stable models. *DNMT3A* and *TET2* are the top two most frequently mutated epigenetic regulators. 2) *DNMT3A* and *TET2* are epigenetic regulator genes, directly impacting the methylation landscape and leaving a distinguishable signature on the DNA when mutated^17,30^. 3) *DNMT3A* and *TET2* are known to confer a broad risk of converting to a myeloid malignancy with the acquisition of a cooperating mutation^12^, making them important players of early AML etiology. 4) Mutations in these genes are found in almost all types of hematologic cancers^31^, broadening the impact of our models.

Frequencies of *DNMT3A* and *TET2* mutations in AML are 22% and 11% respectively from prior work^20^. The BeatAML dataset have similar frequencies: 22% (49) patients carry *DNMT3A* and 17% (37) carry *TET2* mutations.

We trained classifiers to infer the wild-type vs. mutant genotypes for *DNMT3A* and *TET2*. Regression models predict the mutation status of a given frequent epigenetic regulator from the downstream methylation changes. We calculated performance measures using receiver operator characteristic (ROC) curve analysis. Our results show that we trained highly robust and accurate classifiers with AUROC = 0.9 for *DNMT3A* and AUROC = 0.8 for *TET2* **(Figure 2A-B)**. This suggests that despite the complex genetic and epigenetic makeup of these post-diagnosis samples, the distinctive signatures left by epigenetic regulator mutations on DNA can be successfully extracted.

**Figure 2.A-B:**
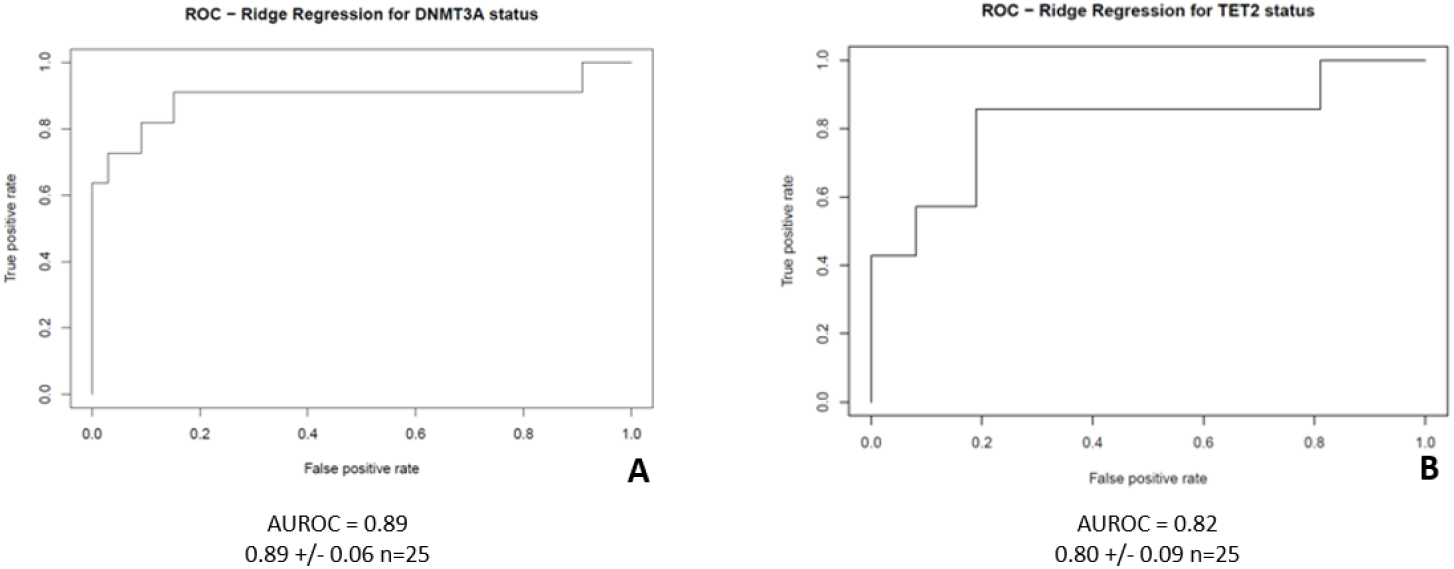
Performance of classifiers measured with AUROC (Area under the ROC curve): Regression models accurately classified wild type vs mutant genotypes based on their down stream impacts.

### DM Rs selected by *DNMT3A* and *TET2* signatures highlight key downstream pathways

To gain more insight into the molecular processes that were indicative of the mutation status of *DNMT3A* and *TET2*, we annotated the genomic sites that has the highest coefficients in the model with the closest gene in the genome **(Tables 1 and 2)**. We found that well-known targets of *DNMT3A*, such as *EVI1/MECOM*^32^ and AEBP2 (a member of the polycomb repressive complex), were often selected by the models. For example, hypomethylation of AEBP2 is strongly associated with *DNMT3A* mutations, and recent single-cell studies have suggested a link between mutated *DNMT3A* and preferential hypomethylation of targets of the polycomb repressive complex 2^33^. Known targets of *TET2* mutations were also selected by the models, including the *CEBPA-AS1* and ABR hypomethylation. Studies have shown that *CEBPA* and *TET2* mutations frequently co-occur and ABR is a transcriptional regulator of CEBPA and contributes to myeloid differentiation. In addition to known targets, we have also identified several novel loci that are strongly correlated with the mutation status of these genes. Further study of these sites could provide new insights into the role of methylation regulation in AML.

**Table 1.** Gene annotations for the top ten most frequently selected features for the *DNM T3A* status prediction both for hypermethylated (green) and hypomethylated (yellow) sites.

| Gene | Potentially Relevant Associations |
| --- | --- |
| *TM2D2* | Found in 8p23.1 amp, linked to pediatric malignancies<sup>38</sup> |
| *HYDIN* | Indicated in familial MDS/AML<sup>39</sup> |
| NA |  |
| *COX7A2* | Part of a myeloid differentiation block regulated by homeobox TFs<sup>40</sup> |
| *SAPCD2P2;VN1R28P* | Pseudogene |
| *WNK2* | MEK pathway upstream regulator<sup>41</sup> |
| *CHN2* | Expression changes associated with lymphomas<sup>42,43</sup> |
| N/A |  |
| N/A |  |
| *TENM2* | Mutually exclusive with *MECOM* and *DNMT3A*, identified in BeatAML<sup>20</sup> |
| N/A |  |
| *HOXB2* | Homeobox TF, known suppressor of *FLT3-ITD* driven AML<sup>44</sup> |
| *C1orf87* |  |
| *DYSF* | Plays a role in monocyte differentiation<sup>45,46</sup> |
| *AEBP2* | Polycomb repressor complex member, frequently deleted in AML, associated with *DNMT3A* hypomethylation<sup>33,47</sup> |
| *VKORC1L1* | Next to *PIK3CG* – a potential tumor suppressor of AML<sup>48,49</sup> |
| *TXNRD1* | Major regulator of metabolism in leukemia cells<sup>50</sup> |
| *AGAP1* | Part of a myeloid self-renewal block regulated by homeobox TFs<sup>51</sup> |
| *RPS6KA2* | rho gtpase<sup>52</sup> |
| *MECOM* | Translocations transform to AML/ mutually exclusive with *TENM2* and *DNMT3A*<sup>32</sup> |

**Table 2.** Gene annotations for the top ten most frequently selected features for the *TET2* status prediction both for hypermethylated (green) and hypomethylated (yellow) sites.

| Gene | Potentially Relevant Associations |
| --- | --- |
| *FRMD6-AS2* |  |
| *FOXK2;RP13-638C3.3* | Genomic stability, DNA repair, cancer stem cell maintenance, cell proliferation, apoptosis and cell metabolism<sup>53</sup> |
| *GTDC1* | Glucosyltransferase high expression in blood leukocytes<sup>54</sup>, 3' MLL fusion partner in acute leukemia<sup>55</sup> |
| *RP11-804A23.1* |  |
| *SCUBE1;Z99756.1* | Initiation and maintenance of MLL-AF9-induced leukemogenesis in vivo. Binds to *FLT3*<sup>56</sup> |
| *TBCD* | Shown to be differentially methylated during granulopoiesis<sup>57</sup> |
| *PHACTR1* | Cytoskeletal regulation<sup>58</sup> |
| *EHD4* |  |
| *DTHD1* | Apoptotic control<sup>59</sup> |
| *GDF7* |  |
| *FAM155A* |  |
| *IQSEC1* |  |
| *TRAP1* | Key mitochondrial regulator<sup>60</sup> |
| *CEBPA-AS1;CTD-2540B15.9* | *CEBPA* regulator, In *CEBPAdm* cases for concomitant mutations, *TET2* found to be most frequently mutated<sup>61</sup> |
| *C1orf112;SELL* |  |
| *ABR* | *CEBPA* regulator, TF contributes to myeloid differentiation<sup>62</sup> |
| *TPO* |  |
| *RP11-191L9.4* |  |
| *RGS17* | *GPCR* regulator<sup>63</sup> |
| *DMRTA1* | Cell differentiation<sup>64</sup> |

### Factorization of methylation changes into Topics link genomic events into distinct methylation programs

Using topic modeling, we reduced the methylation profiles to a latent space of 15 topics. For each patient, topic values represent probabilities of enrichment for each topic based on the most co-variable DMRs across patients. We observe that 15 topics captured major mutations and cytogenetic events as well as having a prior MDS, gender and ELN-2017 risk stratification by genetics^23^. Each patient has different combinations of driver events, but topic modeling successfully deconvoluted the impact of these drivers in the methylome, even for the relatively low frequency chromosomal alterations (**Figure 3**). **Figure 4** tabulates the statistical enrichment of known factors for each topic. For each topic and epigenetic factor pair, we calculated whether that topic’s value is significantly higher in patients with the factor compared to patients without the factor to methodologically quantify enrichment.

**Figure 3:**
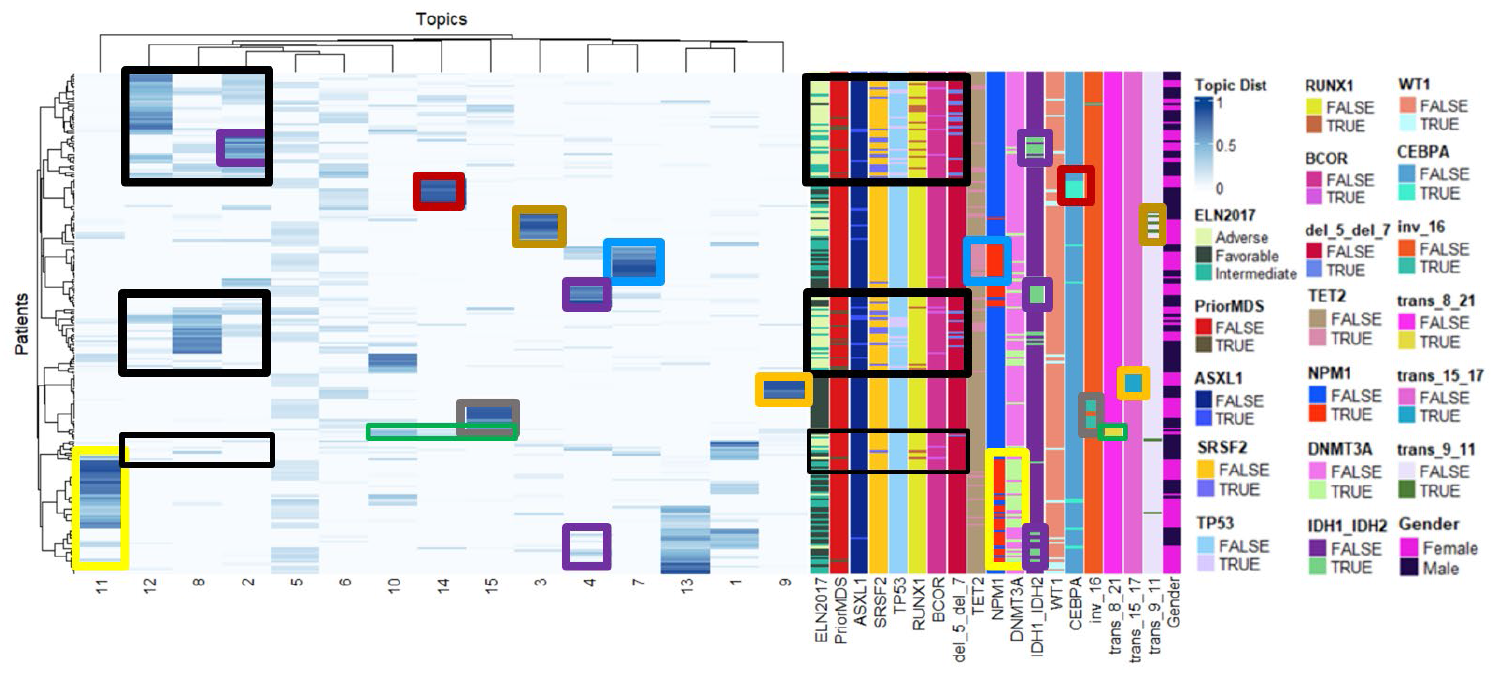
Topic Distribution for 220 post-diagnosis AML patients: Topic size is 15 (columns). For each patient, topic values represent probabilities of enrichment for each topic based on the most variable DMRs across AML patients. Topic 11 is enriched for *DNMT3A* and *NPM1* co-mutations (yellow rectangles). Topic 7 is enriched for *TET2* and *NPM1* co-mutations (blue rectangles). Topic 4 (primarily) and topic 2 are enriched for *IDH1* and *IDH2* (purple rectangles). *NPM1*’s effect is further distributed to topics 4 and 13. Topic 15 is enriched for inv16 (grey rectangles) and t(8,21) and topic 14 is enriched for *CEBPA* (red rectangles) and t(8,21). t(8,21)’s effect is further distributed to topic 10 (green rectangles). Topic 5 is enriched for gender (blue arrows). *ASXL1, SRSF2, TP53, RUNX1, BCOR* and 5q and 7q deletions associated topics are aligning with Prior MDS and ELN-2017 adverse category (black rectangles). The other common chromosomal events t(15, 17) and t(9,11) almost exclusively map to topics 9 and 3 respectively (orange and gold rectangles respectively).

**Figure 4.**
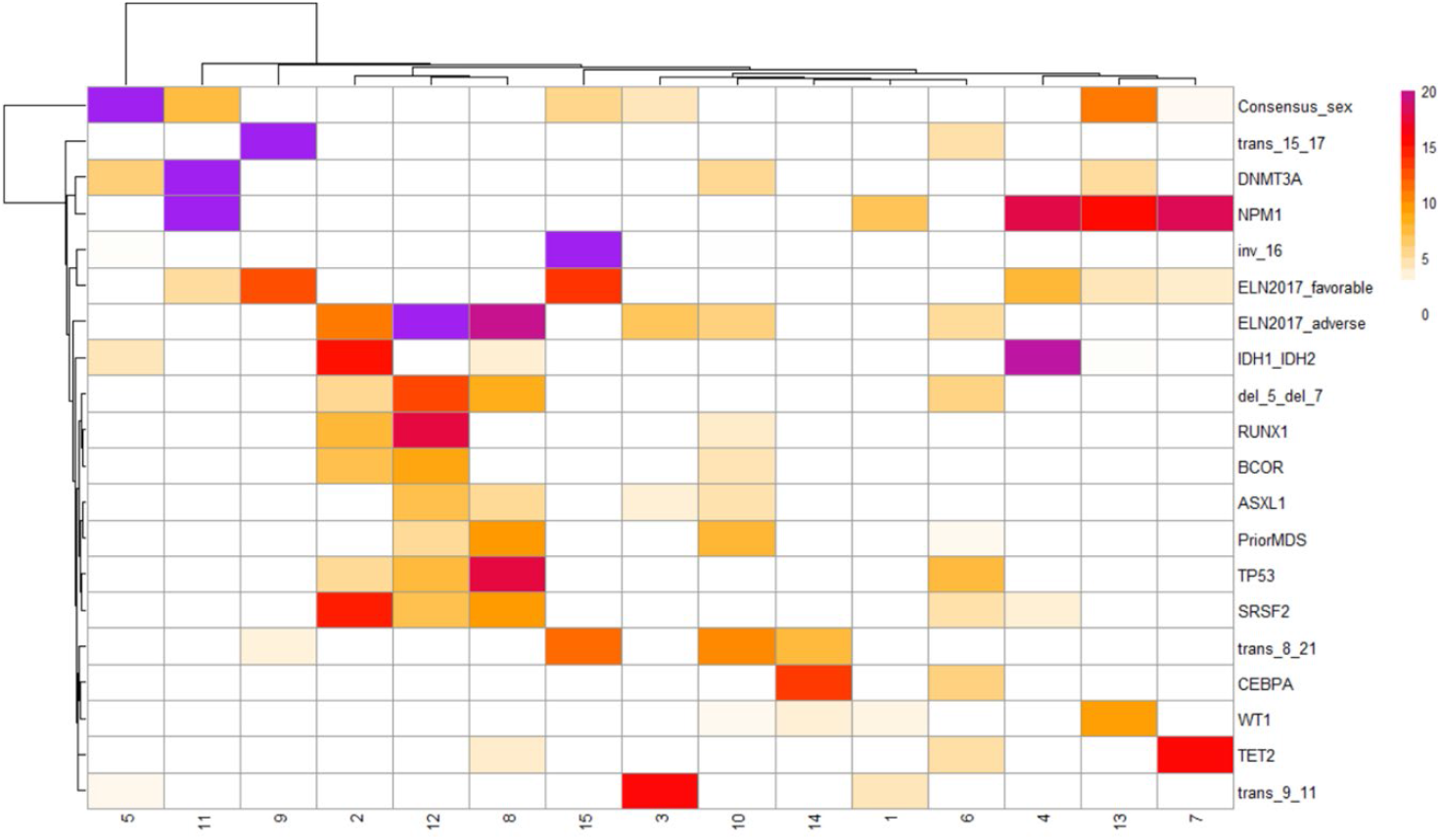
Enrichment heatmap of topics vs. epigenetic factors: Cell colors indicate the significance of whether the average topic values of the patients with the factor (e.g. *DNMT3A* mutant) is significantly higher than the average topic values of the patients without that factor (e.g. *DNMT3A* WT), reflecting –log p-values based on the one sided t-test. Gender is strongly associated with topic 5, *NPM1* and *DNMT3A* co-mutations, topic 11, t(15, 17), topic 9, (inv 16), topic 15, *IDH1* and *IDH2*, topic 4 (primarily) and 2, t(9, 11), topic 3, *CEBPA*, topic 14 (primarily), and *NPM1* and *TET2* co-mutations, topic 7. *NPM1*’s effect is further distributed to topics 4 and 13. t(8,21)’s effect is distributed to topic 15 (primarily), 10 and 14. *RUNX1, BCOR, ASXL1, TP53, SRSF2*, 5q and 7q deletions, having a prior MDS and ELN2017 adverse risk category associated topics are clustered together.

Strong statistical associations between eight topics and specific driving events including topics that represent gender (topic 5) *NPM1* and *DNMT3A* co-mutations (topic 11), translocation of chromosomes 15 and17 (topic 9), inversion of chromosome 16 (topic 15), *IDH1* or *IDH2* mutation (topic 4 and less strongly to topic 2), translocation of chromosomes 9 and 11 (topic 3), *CEBPA* mutation (topic 14) and *NPM1* and *TET2* co-mutations (topic 7) are identified (**Figures 3** and **4**). The effect of *NPM1* is also distributed to topics 4 and 13, while the effect of translocations of chromosome 8 and 21 is distributed to topics 15 (primarily), 10 and 14. Topics associated with *RUNX1, BCOR, ASXL1, TP53, SRSF2*, 5q and 7q deletions, having prior MDS and the ELN2017 adverse risk category are clustered together.

We observe that these epigenetic topics can be broadly classified into four groups for AML **a)** gender associated, **b)** cytogenetic events with a distinct signature, **c)** co-mutations with *NPM1* **d)** A heterogenous set with *TP53, SRSF2, ASXL1, BCOR, RUNX1* mutations and 5q and 7q deletions that is associated with prior MDS and adverse prognosis. As an additional finding, our models show that cytogenetic events, such as t(15;17) have widespread *trans* downstream methylation impacts (as opposed to *cis* aberrations limited to the chromosomes 15 and 17). `

We then asked if the DMRs that have high coefficients for a given topic recapitulate known downstream pathways or reveal novel biology. We annotated each locus with the closest gene in the genome (**Table 3** and **Supplementary Figure 1**). We observe that topic group **a** (1, 5, 6) predictably have DMRs located on chromosomes X and Y. For other topics, multiple downstream pathways have been found to occur repeatedly across various topics, including pathways associated with homeobox (*HOX*) genes, histone deacetylases, lipid metabolism and maintenance of stem-cell state. Topic group **d** (2,8,12) had substantial enrichment in remodeling and differentiation pathways as well as expression of T-Cell receptors, potentially indicating a de-differentiation mechanism to other hematopoietic lineages.

**Table 3.**
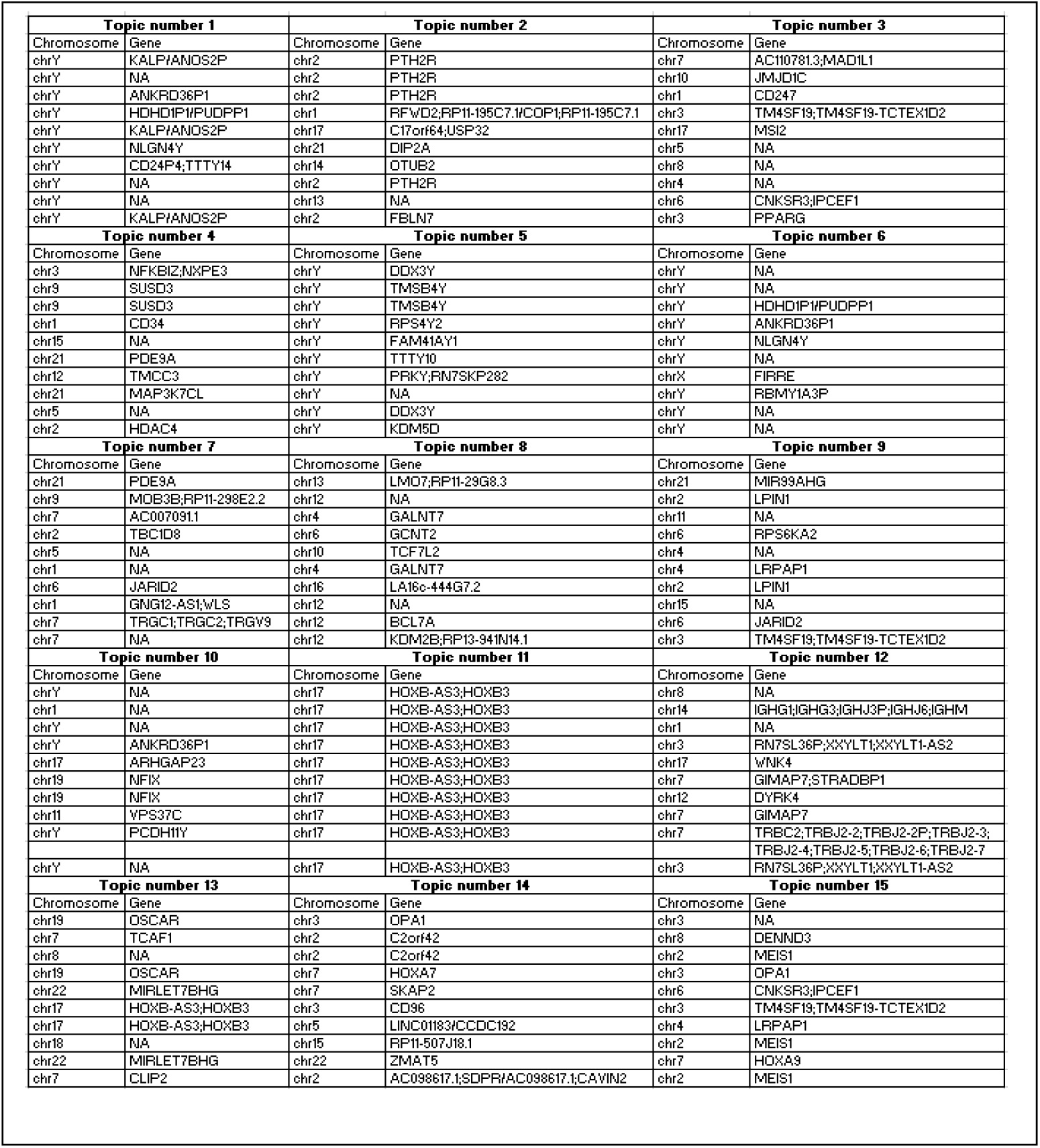
Annotation of the top 10 most enriched loci for all 15 topics.

**Supplementary Figure 2** and **Supplementary Figure 3** show ELN-2017 adverse and ELN-2017 favorable risk category groups of mutations and events clustering together, respectively. In addition, we showed that there is no topic enriched in association with *FLT3* mutations—this is in agreement with previous findings^34^ (**Supplementary Figure 4**).

We also tested whether topic enrichment for cytogenetic events, such as t(15;17) are achieved through their downstream methylation impact or through the detection of their break points by the topic modeling algorithm. For example, topic number 9 is exclusively enriched for t(15; 17) (**Figures 3 and 4**). We annotated the top ten genomic sites with the highest assigned topic values on the genome and none were mapped on chromosomes 15 or 17 (**Table 3 and**

**Supplementary Figure 4**), suggesting that cytogenetic aberrations are categorized based on their *trans*, downstream methylation impact rather than *cis* effects.

## DISCUSSION

We systematically modeled the epigenetic impact of genomic events using supervised and unsupervised approaches. Individual *DNMT3A* and *TET2* signatures were detected with high accuracy and robustness, as they yielded high AUROCs with a testing error that was consistent with training error across multiple training rounds, even amidst the complex genetic and epigenetic landscape of post-diagnosis samples. Methylation loci commonly chosen by the models includes well-known downstream targets such as *EVI1/MECOM* and *AEBP2* for *DNMT3A*, and *CEBPA-AF1* for *TET2*, all of which exhibited hypomethylation. A hypomethylated loci near *HOXB2* was a strong predictor of mutant *DNMT3A* status although there are conflicting reports in the literature of the overall effect of *DNMT3A* mutation on the methylation of the *HOX* promoters^35–37^. This observation strongly suggests that specific loci might have stronger signal for the upstream regulator status as opposed to methylation patterns over a genomic region.

Further investigation of these sites can lead to new mechanistic insight on methylation pathways important in leukemogenesis. As the cohorts become larger, these approaches can be extended beyond *DNMT3A* and *TET2* to other potential AML epigenetic drivers including cytogenetic events and related methylation pathways can be investigated.

Our regression models require labeled data (such as mutation status) and allow us to model the impact of each driving factor (e.g. *DNMT3A* mutation status) independently. However, we need substantially larger cohorts for building models of infrequent drivers such as cytogenetic events. Furthermore, the analysis may be confounded by co-occurring mutations. To complement this approach, we used topic modeling on the same dataset to factorize the methylation profiles.

This yielded a representation of each patient’s methylation pattern as a combination of multiple topics. For each topic we identified we identified mutations and clinical classes that are statistically enriched. We observe that identified topics fell into 4 broad categories: Gender associated, cytogenetic events, *NPM1* co-mutations and a heterogeneous set of mutations in *TP53, SRSF2, ASXL1, BCOR, RUNX1*, and deletions of chromosomes 5q and 7q. The latter group is associated with adverse-risk prognosis.

We have shown that DNA methylation-based categorization achieved by topic modeling resulted in a good concordance with the ELN-2017 risk stratification by genetics and having a prior MDS. Vosberg *et al*.^16^ and Figueroa *et al*^34^ previously pointed to the potential of DNA methylation profiling to refine AML risk stratification and our study reinforces these observations. However, we improve upon these hard-label clustering techniques by representing each patient’s pattern as a combination of several factors as opposed to belonging to a single subtype. This factorization and deconvolution are crucial, given the highly combinatorial nature of the disease. We, in fact see that less frequent factors such as cytogenetic events crosscut previously defined subtypes, methylation effects of which have been obfuscated by more frequent drivers in a hard-clustering approach. Through topic modeling, we can deconvolute the impact of overlapping factors, creating an improved latent representation of the disease state.

As a result of this, our models have made substantial improvements in methylation-based risk-stratification by uncovering previously unknown signatures associated with relatively rare cytogenetic events. This may greatly enhance the subtyping of patients for drug response studies and provide a framework for using methylation as a biomarker in subsequent studies.

We reviewed the frequently selected methylation loci to gain insight on the downstream methylation impact. We observed multiple downstream pathways re-occurring across multiple topics including *HOX* genes, stem-cell and de-differentiation associated pathways as well as histone deacetylases. This implies that although different genetic regulators have distinct methylation impact, they might be converging on common downstream processes and these convergence points might be excellent therapeutic targets. We also observed several downstream pathways that were not canonically associated with upstream genetic events such as parathyroid hormone receptor, lipid homeostasis and protein glycosylation. Perhaps more importantly, we observed that the topic modeling was able to systematically capture the impact of key epigenetic regulating events and simultaneously grouped events that have similar or synergistic impacts together.

As these signatures help us quantify the relationship between a driver event and its downstream methylation impact, they can be leveraged to build predictive models for AML-risk. Methylation changes can be tracked throughout leukemogenesis in early detection cohorts through simple liquid biopsies. Risk-stratification based on downstream epigenetic signatures can help us determine which patients are at risk and who should be monitored more frequently. We can also use these models to test if the function of a regulator is restored or disrupted following treatment, such as hypomethylating agents targeting epigenetic regulator functioning, and stratify who can benefit from the treatment. Since modifications by epigenetic mutations are reversible with therapy, broadening our understanding of the impact of epigenetic mutations on leukemogenesis and therapeutic response will be essential for advancing treatment of myeloid malignancies^31^.

## ACKNOWLEDGEMENT

BG and PTS acknowledges support from the Cancer Early Detection Advanced Research Center (CEDAR) at the Knight Cancer Institute. JWT acknowledges support from the Aqcuired Resistance to Therapy Network (ARTNet), National Institutes of Health (NIH), and National Cancer Institute (NCI) grant U54CA224019. JWT also received support from NCI award R01CA262758 (JWT, SEK), the V Foundation for Cancer Research (JWT), the Gabrielle’s Angel Foundation for Cancer Research (J.W.T.), the Anna Fuller Fund (J.W.T.), the Mark Foundation for Cancer Research (J.W.T.), and the Silver Family Foundation (J.W.T.).

The authors thank the Knight Cancer Institute (NCI P30 CA069533), Bioinformatics, and CEDAR. We are grateful to members of the Spellman Lab, Blundell Lab and to members of the CEDAR; especially to Sebnem Ece Eksi for editing the manuscript and providing her insights, Gurkan Yardimci, Hisham Mohammed, Stefanie Lynch, Ruslan Strogantsev and Jose Luis Montoya Mira for rich discussions.

## AUTHORSHIP CONTRIBUTIONS

BG built the statistical models and methods, analyzed and interpreted the data, plotted the figures. ED contributed to outlining the aim of the project and to our understanding of the computational methods and their applications around biology. JWT built collaborations across different labs, advised BG on the AML aspects of the analyses. BJD provided resources, mentored and advised BG. PTS supervised and mentored BG and reviewed all preceding and analyses herein and provided feedback. All authors read and approved the final manuscript.

## SUPPLEMENTAL DATA

**Supplementary Figure 1.**
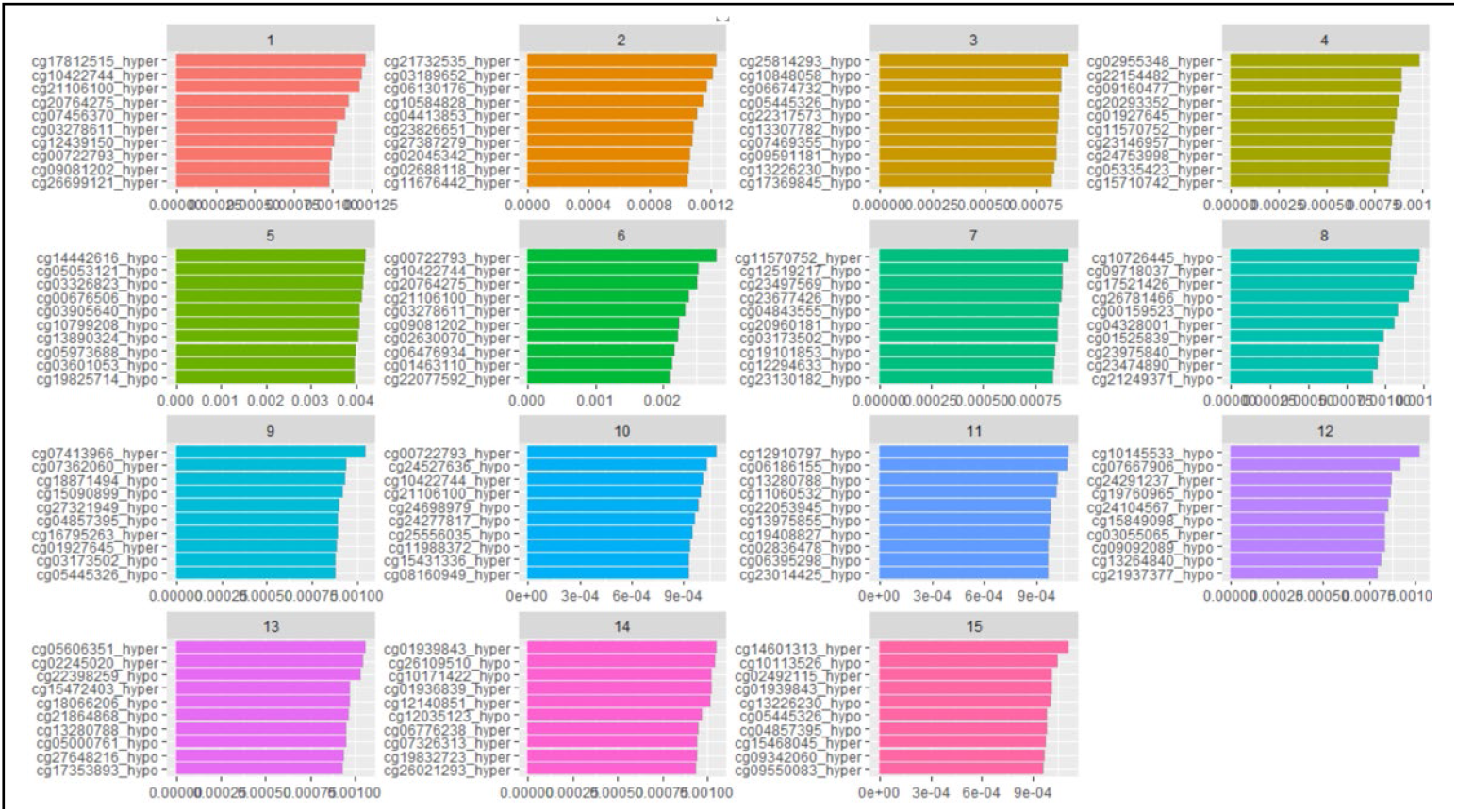
Top 10 most enriched loci for 15 topics: The top 10 probes with the highest assigned topic values and their methylation status (whether they are hyper or hypo methylated) are indicated for all 15 topics. We observe that topics 1, 5 and 6 map to gender and predictably have DMRs located on chromosomes X and Y. Topic 3 is exclusively associated with t(9;11). Notable enriched DMRs are located near: *MSI2*, a translocation partner with *HOXA9, EVI1, TTC40*, and *PAX5* in leukemias^65–68^; *PPARG* – a key regulator for apoptosis and survival and *CNKSR3* – a gene that was identified as a commonly upregulated target in all *AF9/AF10* rearranged AMLs that includes t(9;11)^69^. Topic 9, enriched for t(15; 17) driver events have DMRs near *JARID2*, a known tumor suppressor in AML^70^, *LRPAP1* and *LPIN1*, regulators of lipid hematopoiesis implicated in AML progression^71^. Topic 15, enriched for inv (16) driver events have DMRs around *HOXA9* and *MEIS1*, a frequently implicated pathway in AML progression. Another gene was *OPA1* that is known to be upregulated in AML and a mechanistic component for venetoclax resistance^72^. Topic 11, enriched for co-mutations in *DNMT3A* and *NPM1* has a highly specific signature composed of multiple DMRs centered around *HOXB3* gene – another key homeobox family protein in AML progression^73^. Topic 4 is strongly associated with *NPM1* and *IDH1/IDH2* mutations and interestingly include a DMR close to *CD34*, the definitive marker for hematopoietic stem/progenitor cells. Other AML associated genes selected in Topic 4 include *HDAC4*, a key epigenetic regulator in AML^74,75^. Topic 13, enriched for *NPM1* and *WT1* mutations, also has DMRs near *HOXB3*. Other notable genes include *OSCAR*, a regulator of osteoclast differentiation^76^ and *MIRLET7BHG*, an autophagy regulating lncRNA implicated as an AML survival marker^77^. Topic 7, associated with co-occurring mutations in *NPM1* and *TET2*, has a DMR near *JARID2* a known tumor suppressor in AML^70^, and *TBC1D8*, a known target of *HDAC2* in AML^78^. Topics 2, 8 and 12 have shared components and is associated with ELN adverse category and a complex set of associated mutations in *RUNX1, BCOR, ASXL1, TP53, SRSF2* and 5q and 7q deletions. Topic 2 has multiple DMRs near *PTH2R*, parathyroid hormone receptor, which was shown to be the most upregulated gene in MDS and AML^79^ and differentially expressed in patients with *IDH2* mutations^80^. Topic 8 has DMRs in close proximity to *KDM2B*, a key lysine demethylase in AML; *BCL7A*, a *BAF* remodeling tumor suppressor^81^, and *TCF7L2*, a *WNT* pathway transcription factor implicated in regeneration of hematopoietic stem cells^82^. Topic 12 has DMRs near immunoglobulin heavy constant gamma as well as T-Cell Receptor Beta locus. Other notable genes include *WNK4*^83^, *DYRK1*^84^ and *GIMAP7*^85^ all indicated in stem cell-like signatures. Weak enrichments for *CEBPA* mutations and t(8;21) exist for topics 14 and 10 respectively.

**Supplementary Figure 2.**
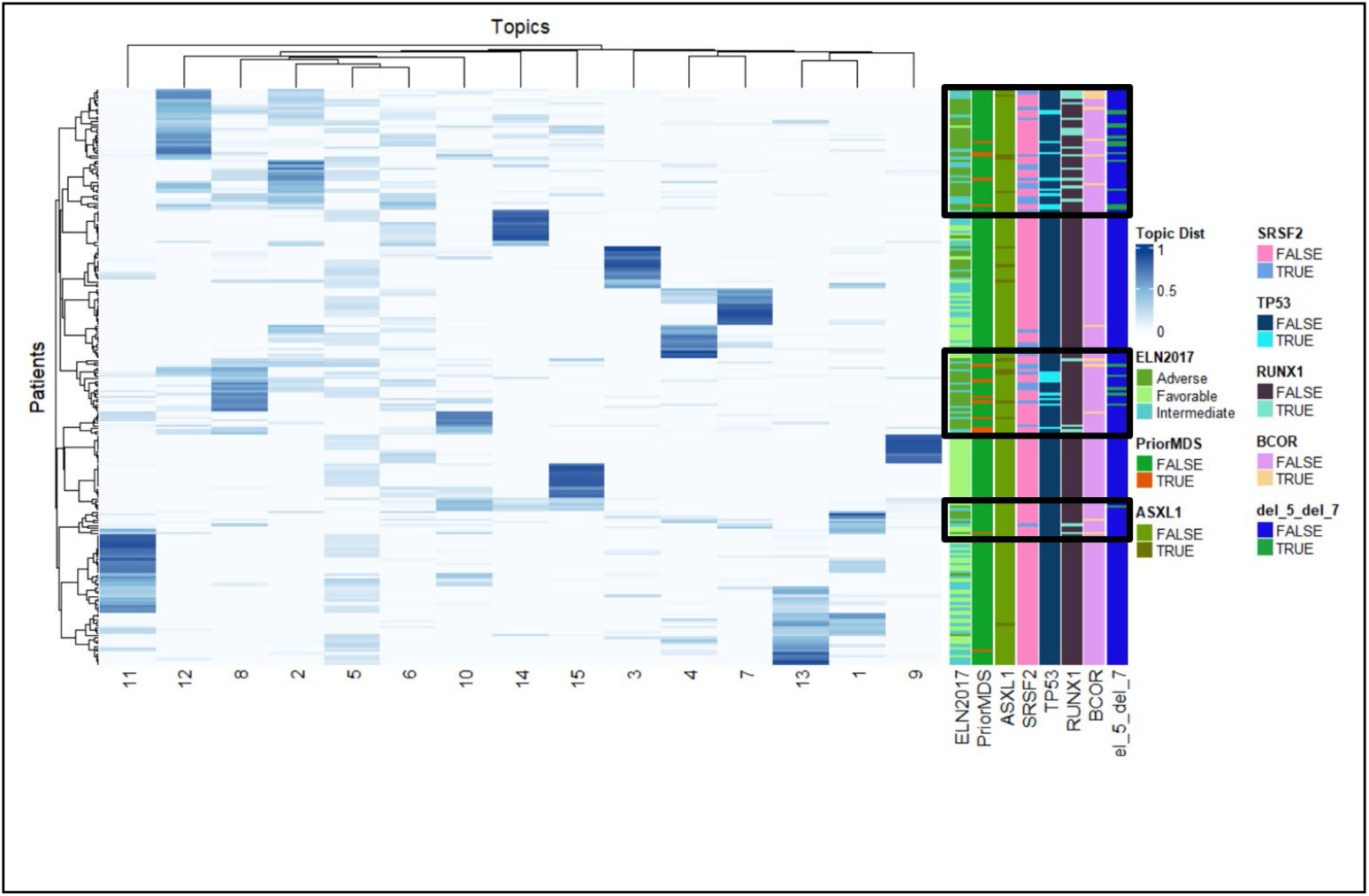
Mutations and events associated with ELN intermediate to adverse risk category: *RUNX1, BCOR, ASXL1, TP53, SRSF2*, 5q and 7q deletions, having a prior MDS and ELN2017 adverse risk category associated topics are clustered together, shown in black rectangles.

**Supplementary Figure 3.**
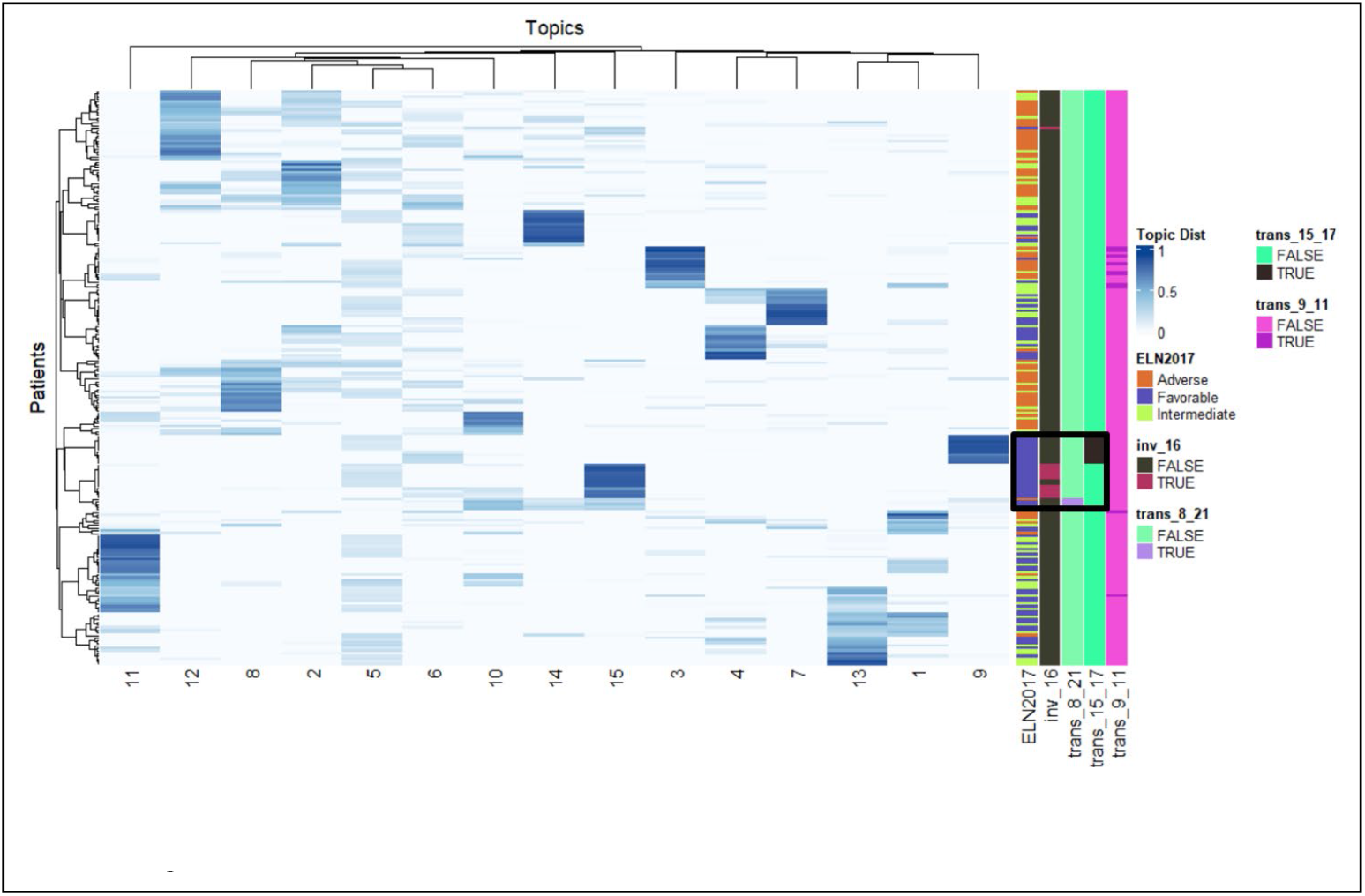
Cytogenetic events associated with ELN favorable risk category: t(15, 17), (inv 16) and t(8, 21) and ELN2017 favorable risk category associated topics are clustered together, shown in the black rectangle.

**Supplementary Figure 4.**
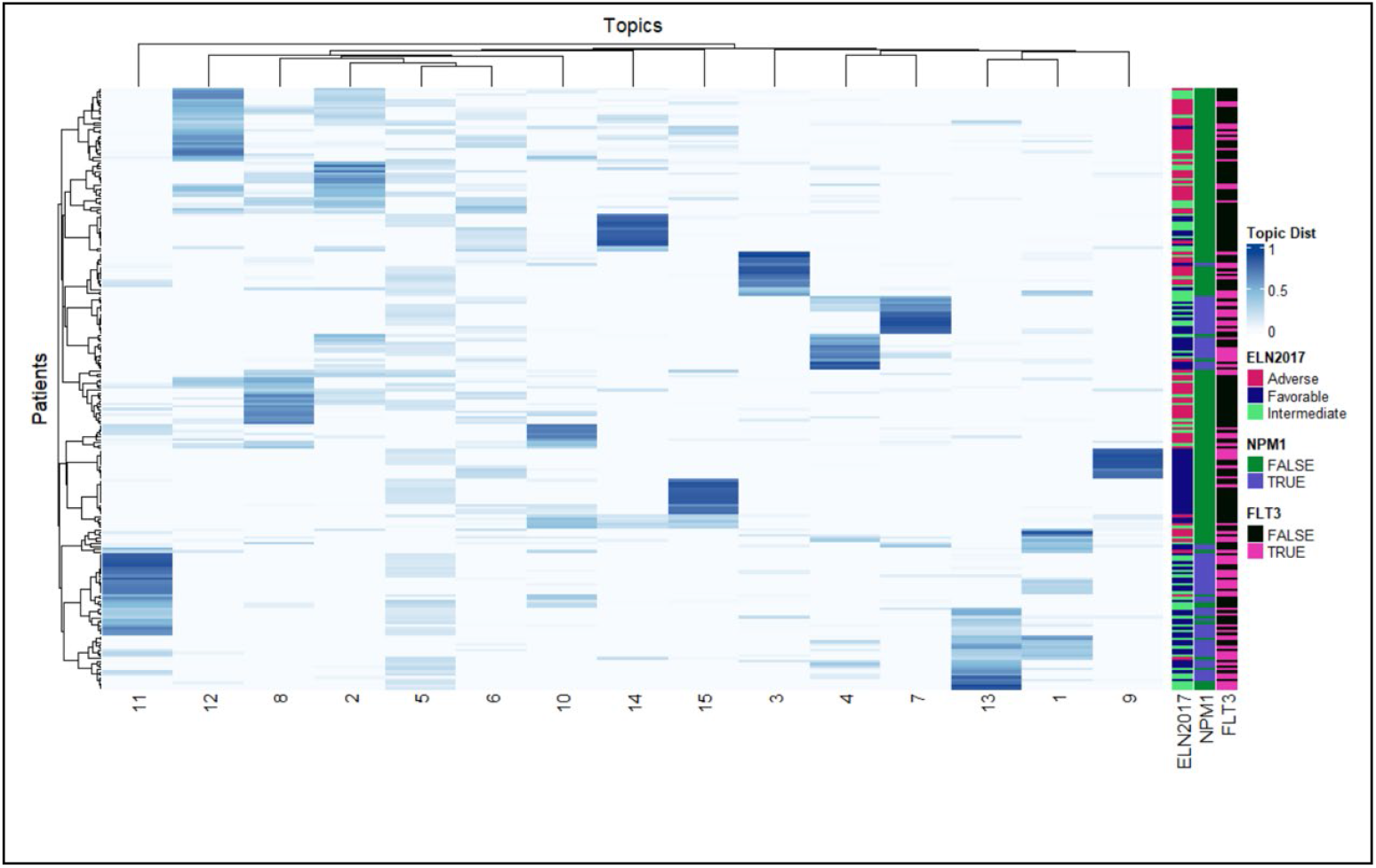
Methylation signature of *FLT3*: *FLT3* mutant samples (in pink) doesn’t have a distinguisahble methylation signature.

